# Influenza sequence validation and annotation using VADR

**DOI:** 10.1101/2024.03.21.585980

**Authors:** Vincent C. Calhoun, Eneida L. Hatcher, Linda Yankie, Eric P. Nawrocki

## Abstract

Tens of thousands of influenza sequences are deposited into the GenBank database each year. The software tool FLAN has been used by GenBank since 2007 to validate and annotate incoming influenza sequence submissions, and has been publicly available as a webserver but not as a standalone tool. VADR is a general sequence validation and annotation software package used by GenBank for Norovirus, Dengue virus and SARS-CoV-2 virus sequence processing that is available as a standalone tool. We have created VADR influenza models based on the FLAN reference sequences and adapted VADR to accurately annotate influenza sequences. VADR and FLAN show consistent results on the vast majority of influenza sequences, and when they disagree VADR is usually correct. VADR can also accurately process influenza D sequences as well as influenza A H17, H18, H19, N10 and N11 subtype sequences, which FLAN cannot. VADR 1.6.3 and the associated influenza models are now freely available for users to download and use.

## Introduction

The World Health Organization estimates that influenza virus infects 1 billion people worldwide each year, leading to between 290,000 and 650,000 deaths [2]. Influenza viruses are segmented negative-sense RNA viruses belonging to the family *Orthomyxoviridae*, of which four genera, commonly referred to as types A, B, C and D, infect vertebrates. The vast majority of human illness from influenza is caused by type A which has caused four pandemics since 1900 (1918, 1957, 1968, 2009) [12, 25] and is also widespread in birds [17] and pigs as well as other mammals. Type B and C also infect humans, and type D is mostly found in cows and pigs. Influenza A has eight segments and is further classified into subtypes (e.g. H5N1) based on the hemagglutinin and neuraminidase glycoproteins encoded on segments 4 and 6 respectively.

Large-scale genomic sequencing of influenza has been employed for nearly twenty years to help understand and track the prevalence, evolution and antiviral resistance of the virus [9, 22, 28, 14]. Nearly one million genomic (nucleotide) influenza sequences exist in the public databases GenBank, European Nucleotide Archive (ENA) [6], and DNA Databank of Japan (DDBJ) [24] which comprise the International Nucleotide Sequence Database Collaboration (INSDC) [4]. While all three INSDC databases share and host the same data, sequences are initially submitted to, and quality checked by, only one of the databases. More than 90% of the influenza sequences in INSDC databases were submitted to GenBank, which is maintained by the National Center for Biotechnology Information (NCBI) at the National Library of Medicine (NLM) in the United States (see Table 1). Since 2018, roughly 50,000 influenza A sequences per year were deposited in GenBank. The volume of influenza B sequences has decreased in recent years, and the volume of influenza C and D sequences has always been relatively low, never reaching 1000 in a year. In addition to hosting the sequence information, NCBI provides the NCBI Virus resource [10] to facilitate and simplify access to it.

**Table 1.** Number of influenza virus sequences deposited in GenBank database since 2018. Sequence counts were obtained using the NCBI Virus Data Hub filtering by release date. “all” row includes total counts of all sequences (including before 2018). “GenBank” row indicates the fraction of all influenza sequences in INSDC databases that were submitted to GenBank (not to EMBL or DDBJ). NCBI taxonomy ids: 11320 (influenza A); 11520 (influenza B); 11552 (influenza C); 1511084 (influenza D).

| year | influenza A | B | C | D |
| --- | --- | --- | --- | --- |
| 2018 | 58719 | 14256 | 23 | 23 |
| 2019 | 100678 | 11622 | 13 | 211 |
| 2020 | 64615 | 13589 | 28 | 436 |
| 2021 | 29021 | 1336 | 133 | 203 |
| 2022 | 73205 | 1375 | 39 | 305 |
| 2023 | 103869 | 7325 | 36 | 19 |
| all | 940860 | 126333 | 525 | 1409 |
| GenBank | 0.938 | 0.964 | 0.200 | 0.949 |

### FLAN is a tool for influenza genome annotation

Since 2007, NCBI has used an internally developed software program called FLAN (FLu ANnotation tool) [5] for validation and annotation of influenza sequences. FLAN has been used in two main contexts: for screening incoming influenza sequence submissions to the GenBank database, and as a publicly available webserver that allows users to validate and annotate their own data. As a screening tool for GenBank, FLAN has been used since 2017 to automatically process sequence submissions of influenza A, B, or C sequences. Submissions with zero FLAN errors are automatically deposited into GenBank without any manual processing. Submissions with at least one sequence with one or more errors are not deposited. Instead, detailed error reports are either automatically sent to the submitters or reviewed by expert NCBI curators for quality, depending on the specific nature of the errors. Prior to 2017, FLAN was used manually to check influenza sequences submitted to GenBank.

FLAN proceeds through multiple steps to classify, validate and annotate input sequences. First, the *blastn* program from the BLAST package [3] is used to compare the input nucleotide sequence against a reference database of influenza A, B, and C nucleotide sequences and classify each sequence type (A, B, or C) and segment. For influenza A segment 4 and 6 sequences, the subtype of the hemagglutinin and neuraminidase segments is also determined. The FLAN reference database contains a single reference nucleotide sequence for each type and segment, and subtype for hemagglutinin (H1 to H16) and neuraminidase (N1 to N10) influenza A segments.

Following classification, each sequence is then aligned to the corresponding reference protein set of one or more proteins for the classified type/segment/subtype using the ProSplign program for nucleotide to protein alignment [1]. ProSplign is capable of detecting frameshifts and handling introns. FLAN detects a dozen types of errors in input sequences listed in Table 2. An error-free alignment is one that extends to the N and C termini of the reference protein sequence (or to the end of the input sequence if it is incomplete at the 5’ and/or 3’ end) with valid start and stop codons at the ends and zero in-frame internal stop codons. Further, there must be zero frameshifts and for proteins with mature peptides those peptides must be properly arranged (adjacent peptides must have zero nucleotides between them and not overlap). FLAN uses the positional information from the ProSplign alignment to determine nucleotide boundary positions for coding sequences and signal and mature peptides for type A segment 4 sequences.

**Table 2.**
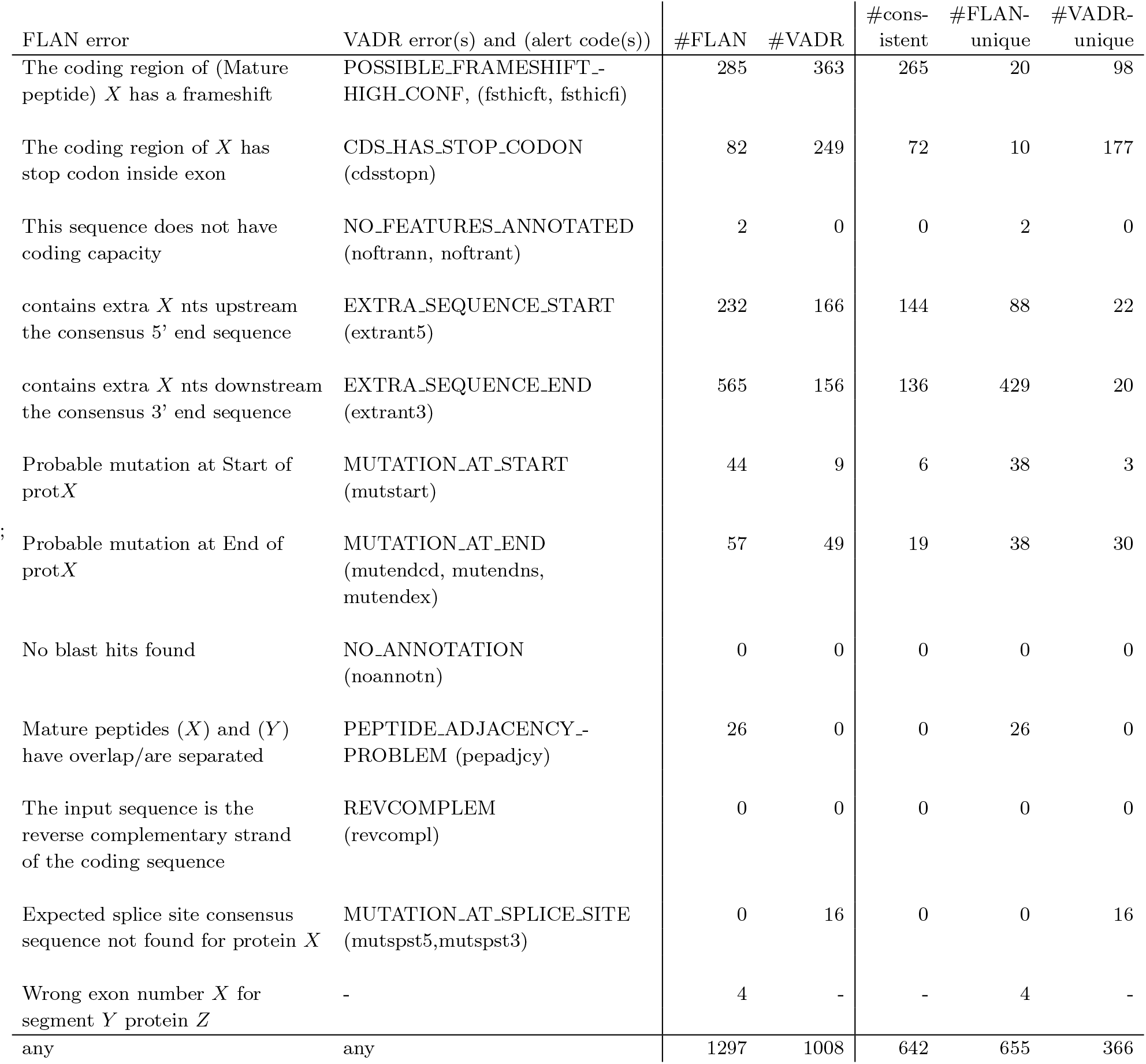
Mapping of FLAN and VADR errors and number of instances in the combined training and testing datasets. Counts are of number of sequence/feature pairs with at least one of the FLAN or VADR error/alert. Some sequence/feature pairs may have multiple errors for the same feature but these are only counted once. Any mapped FLAN and VADR errors are considered consistent if they occur for the same sequence/feature pair. All FLAN errors that cause a sequence to fail (with exceptions detailed in the text) are listed, but not all VADR fatal alerts are.

Although FLAN has been in routine use at NCBI for more than 15 years, it is difficult to maintain and expand for novel influenza sequence diversity. Additionally, the program is not portable or available as a standalone program and users may only access it via its webserver interface or by submitting sequences to GenBank, such that local execution is not possible and high-throughput use is difficult.

### VADR is a general tool for viral genome annotation

VADR (Viral Annotation DefineR) is a software package also developed at NCBI for validating and annotating viral sequences and protein-coding sequences [23]. It is written in Perl but also relies on and calls programs from other software such as Infernal [19], BLAST [3], FASTA [21], and minimap2 [15, 16]. VADR is a general tool that includes a program called *v-build*.*pl* for creating new models for a virus based on existing GenBank or RefSeq records that include information on CDS, gene, mature peptide and structural RNA features. The *v-annotate*.*pl* program then uses a library of those models to classify input sequences to their best-matching model, and validate and annotate features based on an alignment to that model. Finally, the protein coding potential of each predicted coding sequence is validated using *blastx* with a library of reference protein sequences. During this analysis, different types of unexpected features, such as early stop codons and potential frameshifts identified from the nucleotide alignment (not at the protein level) can be identified and reported in the form of *alerts* to the user. A subset of alerts are *fatal* and cause a sequence to *fail* validation.

VADR is used to automatically screen and validate incoming GenBank submissions of Norovirus, Dengue virus, SARS-CoV-2 and metazoan COX1 protein coding mitochondrial sequences in a similar way to how FLAN is used for influenza sequence submissions. Sequence submissions with zero fatal alerts are automatically deposited into GenBank. For submissions with one or more fatal alerts, a detailed error report is sent to the submitter or prepared for a GenBank curator to review. The scope of VADR alerts overlaps heavily with that of FLAN errors (Table 2).

In contrast to FLAN, VADR is actively maintained and developed and is available as a standalone program so that users can run it locally. We describe here our effort to make VADR useful for influenza sequence analysis, by building new models based on the FLAN reference sequences.

## Materials and methods

### Creation of VADR influenza model libraries

We created influenza models for VADR based on the existing FLAN models and then compared the performance of VADR and FLAN at validating and annotating existing influenza sequences. The nucleotide and protein reference sequence sets that FLAN uses are available online at (https://ftp.ncbi.nlm.nih.gov/genomes/INFLUENZA/ANNOTATION/) but the specific accessions of those sequences are absent from those files and from the FLAN publication [5]. The first step towards making VADR influenza models was determining the INSDC accessions that the FLAN reference sequences map to, so that we could build VADR models from those accessions and those models would include the sequence annotation information from GenBank, ENA or DDBJ.

The FLAN reference sequence set includes 46 nucleotide sequences, 96 protein sequences, and 72 mature peptide sequences. Of the 46 nucleotide sequences, 45 have at least one identically matching sequence in INSDC, and the remaining sequence is a subsequence of an existing INSDC sequence. Of the 96 protein sequences, 69 matched identically to at least one INSDC sequence, and 14 of the remaining 28 are an exact subsequence of at least one INSDC sequence. For the remaining 14, we determined the best matching INSDC sequence for each, defined as the sequence with the best *blastp* [3] score when searched against the non-redundant protein sequences set (“nr”) database using the *blastp* webserver (https://blast.ncbi.nlm.nih.gov/Blast.cgi). The 72 mature peptide sequences are not relevant to VADR model building, so we did not attempt to map these to INSDC sequences. More details on the sequence mapping, including specific accessions, can be found in the freely available VADR influenza model dataset.

A VADR model was created for each of the 46 INSDC-mapped nucleotide sequences using the *v-build*.*pl* program from VADR 1.6.3 with default parameters, and gene and CDS product names were modified to match FLAN. The models were combined to create a model library as explained in the VADR documentation. The VADR model library was modified by adding the mapped INSDC protein sequences to their corresponding models’ *blastx* protein libraries.

After building VADR models that matched the FLAN reference sets, we expanded them in several ways in an effort to improve their performance. As detailed below, we added eight additional protein sequences to the influenza A *blastx* libraries. We also added models for influenza D and for influenza A subtypes that were discovered after FLAN was developed, and replaced five incomplete influenza C genome sequences used by FLAN with complete genome sequences from RefSeq. Specifically, we:

- added 8 models for the novel H17/N10 subtypes [26], one per segment (accessions CY103881-CY103888)
- added 8 models for the novel H18/N11 subtypes [27], one per segment (accessions CY125942-CY125949)
- added 1 model for the novel H19 subtype [8], segment 4 (accession ON637239)
- replaced the 5 non-RefSeq influenza C models with RefSeq-based models (GN364866, GM968019, GM968018, GM968017, and GM968016 replaced with NC_006307, NC_006308, NC_006309, NC_006310, and NC_006312, respectively).
- added 7 RefSeq-based influenza D models (NC_036615-NC_036621)

For the 29 added or replaced models, the *v-build*.*pl* program from VADR 1.6.3 was used to create the models with default parameters using the indicated accessions.

We made changes to two of the VADR models to improve annotation on some sequences. Specifically, we rebuilt the covariance model (CM) file for the CY006079 type A segment 5 model using a two sequence alignment containing the original CY006079.1 sequence and a copy of it with an insertion of a single *A* nucleotide after position 1542. Similarly, we rebuilt the CM file for the CY005970 type A segment 4 model using a two sequence alignment containing the original CY005970.1 sequence and a copy of it with a substitution of the *G* nucleotide at position 1719 with an *A*. Using these new models reduces the number of spurious alerts related to stop codons. The changes were motivated by analysis of the VADR results on the training set sequences and more details are available in the documentation included with the model files. Tables 3 and 4 include more details on the final set of VADR influenza models (version 1.6.3-2).

**Table 3.** List of VADR and FLAN influenza A model reference sequences and attributes of associated proteins. The “#coords” column indicates the number of distinct pairs of start and stop genome nucleotide coordinates for all proteins in the set. The “# proteins” column indicates the number of proteins in the set, and the “#extra” column indicates the number of additional proteins added to the VADR protein set not present in the FLAN set based on analysis of training set results. For CDS that have italicized “CDS product” and “gene” names, FLAN errors are converted to warnings and VADR converts them to “misc feature” features if they have certain usually fatal alerts, instead of failing the sequence. Bold model accessions indicate models without an analog in FLAN.

| type | seg | sub-type | model accession | model length | CDS product | gene | intron? | #proteins | #coords | #extra |
| --- | --- | --- | --- | --- | --- | --- | --- | --- | --- | --- |
| A | 1 | - | CY002079 | 2341 | polymerase PB2 | PB2 | no | 3 | 2 | - |
| A | 1 | - | **CY103881** | 2338 | polymerase PB2 | PB2 | no | 1 | 1 | - |
| A | 1 | - | **CY125942** | 2338 | polymerase PB2 | PB2 | no | 1 | 1 | - |
| A | 2 | - | CY003646 | 2341 | *PB1-F2 protein* | *PB1-F2* | no | 14 | 12 | - |
|  |  |  |  |  | polymerase PB1 | PB1 | no | 4 | 2 | 1 |
| A | 2 | - | **CY103882** | 2339 | polymerase PB1 | PB1 | no | 1 | 1 | - |
| A | 2 | - | **CY125943** | 2339 | polymerase PB1 | PB1 | no | 1 | 1 | - |
| A | 3 | - | CY003645 | 2233 | *PA-X protein* | *PA-X* | yes | 4 | 2 | - |
|  |  |  |  |  | polymerase PA | PA | no | 3 | 1 | 1 |
| A | 3 | - | **CY103883** | 2216 | polymerase PA | PA | no | 1 | 1 | - |
| A | 3 | - | **CY125944** | 2216 | polymerase PA | PA | no | 1 | 1 | - |
| A | 4 | H1 | CY000449 | 1778 | hemagglutinin | HA | no | 2 | 1 | - |
| A | 4 | H2 | CY003907 | 1773 | hemagglutinin | HA | no | 2 | 1 | - |
| A | 4 | H3 | CY002000 | 1762 | hemagglutinin | HA | no | 4 | 1 | - |
| A | 4 | H4 | CY004847 | 1738 | hemagglutinin | HA | no | 2 | 1 | - |
| A | 4 | H5 | DQ864721 | 1780 | hemagglutinin | HA | no | 4 | 1 | - |
| A | 4 | H6 | DQ376635 | 1747 | hemagglutinin | HA | no | 2 | 1 | - |
| A | 4 | H7 | CY006037 | 1732 | hemagglutinin | HA | no | 7 | 1 | - |
| A | 4 | H8 | CY005970 | 1744 | hemagglutinin | HA | no | 2 | 1 | - |
| A | 4 | H9 | CY004642 | 1742 | hemagglutinin | HA | no | 3 | 1 | 1 |
| A | 4 | H10 | CY006001 | 1728 | hemagglutinin | HA | no | 2 | 1 | - |
| A | 4 | H11 | CY006005 | 1760 | hemagglutinin | HA | no | 2 | 1 | - |
| A | 4 | H12 | CY006008 | 1737 | hemagglutinin | HA | no | 2 | 1 | - |
| A | 4 | H13 | CY005979 | 1768 | hemagglutinin | HA | no | 2 | 1 | - |
| A | 4 | H14 | M35997 | 1749 | hemagglutinin | HA | no | 1 | 1 | - |
| A | 4 | H15 | CY006034 | 1763 | hemagglutinin | HA | no | 1 | 1 | - |
| A | 4 | H16 | AY684891 | 1760 | hemagglutinin | HA | no | 1 | 1 | - |
| A | 4 | H17 | **CY103884** | 1784 | hemagglutinin | HA | no | 1 | 1 | - |
| A | 4 | H18 | **CY125945** | 1771 | hemagglutinin | HA | no | 1 | 1 | - |
| A | 4 | H19 | **ON637239** | 1686 | hemagglutinin | HA | no | 1 | 1 | - |
| A | 5 | - | CY006079 | 1565 | nucleocapsid protein | NP | no | 2 | 1 | - |
| A | 5 | - | **CY103885** | 1558 | nucleocapsid protein | NP | no | 1 | 1 | - |
| A | 5 | - | **CY125946** | 1557 | nucleocapsid protein | NP | no | 1 | 1 | - |
| A | 6 | N1 | CY002538 | 1463 | neuraminidase | NA | no | 3 | 1 | 1 |
| A | 6 | N2 | CY002010 | 1467 | neuraminidase | NA | no | 4 | 2 | 1 |
| A | 6 | N3 | CY005890 | 1453 | neuraminidase | NA | no | 2 | 1 | - |
| A | 6 | N4 | CY005359 | 1463 | neuraminidase | NA | no | 1 | 1 | - |
| A | 6 | N5 | CY004429 | 1470 | neuraminidase | NA | no | 2 | 1 | - |
| A | 6 | N6 | CY005641 | 1465 | neuraminidase | NA | no | 2 | 1 | - |
| A | 6 | N7 | CY004435 | 1462 | neuraminidase | NA | no | 1 | 1 | - |
| A | 6 | N8 | CY004056 | 1461 | neuraminidase | NA | no | 1 | 1 | - |
| A | 6 | N9 | CY004131 | 1460 | neuraminidase | NA | no | 2 | 1 | - |
| A | 6 | N10 | **CY103886** | 1390 | neuraminidase | NA | no | 1 | 1 | - |
| A | 6 | N11 | **CY125947** | 1426 | neuraminidase-like protein | NA | no | 1 | 1 | - |
| A | 7 | - | CY002009 | 1027 | matrix protein 1 | M1 | no | 2 | 1 | - |
|  |  |  |  |  | matrix protein 2 | M2 | yes | 3 | 2 | - |
| A | 7 | - | **CY103887** | 1027 | matrix protein 1 | M1 | no | 1 | 1 | - |
|  |  |  |  |  | matrix protein 2 | M2 | yes | 1 | 1 | - |
| A | 7 | - | **CY125948** | 1027 | matrix protein 1 | M1 | no | 1 | 1 | - |
|  |  |  |  |  | matrix protein 2 | M2 | yes | 1 | 1 | - |
| A | 8 | - | CY002284 | 890 | nonstructural protein 1 | NS1 | no | 14 | 9 | 3 |
|  |  |  |  |  | nuclear export protein | NEP | yes | 2 | 1 | - |
| A | 8 | - | **CY103888** | 895 | nonstructural protein 1 | NS1 | no | 1 | 1 | - |
|  |  |  |  |  | nuclear export protein | NEP | yes | 1 | 1 | - |
| A | 8 | - | **CY125949** | 895 | nonstructural protein 1 | NS1 | no | 1 | 1 | - |
|  |  |  |  |  | nuclear export protein | NEP | yes | 1 | 1 | - |

**Table 4.** List of VADR and FLAN influenza B, C, and D model reference sequences and attributes of associated proteins. The “#coords” column indicates the number of distinct pairs of start and stop genome nucleotide coordinates for all proteins in the set. The “# proteins” column indicates the number of proteins in the set. For CDS that have italicized “CDS product” and “gene” names, FLAN errors are converted to warnings and VADR converts them to “misc feature” features if they have certain usually fatal alerts, instead of failing the sequence. Bold model accessions indicate models without an analog in FLAN. Italicized model accessions indicate models for which VADR uses a different accession than FLAN.

| type | seg | model<br>accession | model<br>length | CDS product | gene | intron? | #proteins | #coords |
| --- | --- | --- | --- | --- | --- | --- | --- | --- |
| B | 1 | EF626642 | 2369 | polymerase PB1 | PB1 | no | 2 | 1 |
| B | 2 | AY504599 | 2396 | polymerase PB2 | PB2 | no | 2 | 1 |
| B | 3 | EF626633 | 2305 | polymerase PA | PA | no | 2 | 1 |
| B | 4 | AF387493 | 1882 | hemagglutinin | HA | no | 4 | 1 |
| B | 5 | EF626631 | 1844 | nucleoprotein | NP | no | 2 | 1 |
| B | 6 | AY191501 | 1557 | *NB protein* | *NB* | no | 1 | 1 |
|  |  |  |  | neuraminidase | NA | no | 3 | 1 |
| B | 7 | AY504605 | 1190 | BM2 protein | BM2 | no | 4 | 2 |
|  |  |  |  | matrix protein 1 | M1 | no | 1 | 1 |
| B | 8 | AY504614 | 1097 | nonstructural protein 1 | NS1 | no | 4 | 1 |
|  |  |  |  | nuclear export protein | NEP | yes | 3 | 1 |
| C | 1 | *NC_006307* | 2365 | polymerase PB2 | PB2 | no | 1 | 1 |
| C | 2 | *NC_006308* | 2363 | polymerase PB1 | PB1 | no | 1 | 1 |
| C | 3 | *NC_006309* | 2183 | polymerase P3 | P3 | no | 1 | 1 |
| C | 4 | *NC_006310* | 2073 | hemagglutinin-esterase | HE | no | 1 | 1 |
| C | 5 | NC_006311 | 1807 | nucleoprotein | NP | no | 1 | 1 |
| C | 6 | *NC_006312* | 1180 | CM2 protein | CM2 | no | 1 | 1 |
|  |  |  |  | matrix protein 1 | M1 | yes | 1 | 1 |
| C | 7 | NC_006306 | 935 | nonstructural protein 1 | NS1 | no | 2 | 1 |
|  |  |  |  | nonstructural protein 2 | NEP | yes | 2 | 1 |
| D | 1 | **NC.036616** | 2364 | polymerase PB2 | PB2 | no | 1 | 1 |
| D | 2 | **NC.036615** | 2330 | polymerase PB1 | PB1 | no | 1 | 1 |
| D | 3 | **NC.036619** | 2195 | polymerase 3 | P3 | no | 1 | 1 |
| D | 4 | **NC.036618** | 2049 | hemagglutinin-esterase precursor | HEF | no | 1 | 1 |
| D | 5 | **NC.036617** | 1775 | nucleoprotein | NP | no | 1 | 1 |
| D | 6 | **NC.036620** | 1219 | P42 | P42 | no | 1 | 1 |
| D | 7 | **NC.036621** | 868 | nonstructural protein 1 | NS1 | no | 1 | 1 |
|  |  |  |  | nonstructural protein 2 | NS2 | yes | 1 | 1 |

### Construction of training and testing sequence sets

To compare the performance of VADR and FLAN at validating and annotating influenza sequences, we constructed multiple disjoint sequence sets for training and testing. The training sets are made up of 10,000 influenza A sequences, 1000 influenza B sequences, and 500 influenza C sequences each from GenBank and from EMBL or DDBJ (23,000 sequences in total, Table 5), which are 60 nucleotides (nt) or longer and are not in the “patent” INSDC division. Separate sets were chosen from GenBank and EMBL/DDBJ because the vast majority of GenBank influenza sequences submitted since 2007 have been screened with FLAN, whereas EMBL and DDBJ do not use FLAN. The sequences were chosen randomly from all candidate sequences downloaded from the NCBI virus resource (https://www.ncbi.nlm.nih.gov/labs/virus/vssi/#/) with release date before March 17, 2023 (the date of our initial collection). We examined and compared FLAN and VADR results on the training sequences to help us improve VADR performance.

**Table 5.** Comparison of pass/fail outcomes for FLAN and VADR on the influenza sequence training datasets.

| dataset | num seqs | VADR<br>model | fraction |  | #fail both | #FLAN-pass | #FLAN-fail |
| --- | --- | --- | --- | --- | --- | --- | --- |
|  |  |  | pass both | #pass both |  | VADR-fail | VADR-pass |
| flu A GenBank | 10000 | final | 0.989 | 9891 | 54 | 50 | 5 |
|  |  | FLAN | 0.986 | 9859 | 54 | 82 | 5 |
|  |  | FLAN-ntonly | 0.895 | 8951 | 58 | 990 | 1 |
| flu A EMBL+DDBJ | 10000 | final | 0.978 | 9784 | 156 | 50 | 10 |
|  |  | FLAN | 0.973 | 9726 | 156 | 108 | 10 |
|  |  | FLAN-ntonly | 0.923 | 9230 | 156 | 604 | 10 |
| flu B GenBank | 1000 | final | 0.999 | 999 | 0 | 0 | 1 |
|  |  | FLAN | 0.999 | 999 | 0 | 0 | 1 |
|  |  | FLAN-ntonly | 0.944 | 944 | 0 | 55 | 1 |
| flu B EMBL+DDBJ | 1000 | final | 0.981 | 981 | 4 | 1 | 14 |
|  |  | FLAN | 0.981 | 981 | 4 | 1 | 14 |
|  |  | FLAN-ntonly | 0.974 | 974 | 4 | 8 | 14 |
| flu C GenBank | 500 | final | 0.818 | 409 | 1 | 0 | 90 |
|  |  | FLAN | 0.818 | 409 | 90 | 0 | 1 |
|  |  | FLAN-ntonly | 0.818 | 409 | 90 | 0 | 1 |
| flu C EMBL+DDBJ | 500 | final | 0.720 | 360 | 0 | 0 | 140 |
|  |  | FLAN | 0.720 | 360 | 136 | 0 | 4 |
|  |  | FLAN-ntonly | 0.720 | 360 | 136 | 0 | 4 |
| total | 23000 | final | 0.975 | 22424 | 215 | 101 | 260 |
|  |  | FLAN | 0.971 | 22334 | 440 | 191 | 35 |
|  |  | FLAN-ntonly | 0.907 | 20868 | 444 | 1657 | 31 |

We constructed additional sets of testing sequences that we did not utilize until after the VADR improvements based on the training set analysis were complete, to see if the VADR performance on the training sets extended to other sequences. The test sets were constructed similarly to the training sets, again both from GenBank and EMBL or DDBJ but also from two date ranges: up to March 17, 2023, the date of initial collection of the training sets, and between March 18, 2023 and November 30, 2023. Testing on the more current sequences checked performance on potentially novel sequence diversity not present in the other sequences. In total there were 12 sets of test sequences, one for each combination of the three possible influenza types, two possible databases, and two possible date ranges, with a combined 35,555 sequences. The specific numbers in each of the 12 sets is shown in Table 6. Some test sets were smaller than the corresponding training sets if not enough qualifying sequences existed to match the training set size (e.g. only 15 GenBank current influenza C test sequences compared with 500 GenBank influenza C training sequences). The test sequences were constructed such that there were zero sequences in common with the training sets.

**Table 6.** Comparison of pass/fail outcomes for FLAN and VADR on the influenza sequence testing datasets.

| dataset | release date | num seqs | fraction |  | #fail both | #FLAN-pass | #FLAN-fail |
| --- | --- | --- | --- | --- | --- | --- | --- |
|  |  |  | pass both | #pass both |  | VADR-fail | VADR-pass |
| flu A GenBank | before 3/18/2023 | 10000 | 0.992 | 9915 | 50 | 31 | 4 |
|  | after 3/17/2023 | 10000 | 0.998 | 9983 | 0 | 17 | 0 |
| flu A EMBL+DDBJ | before 3/18/2023 | 10000 | 0.979 | 9793 | 156 | 46 | 5 |
|  | after 3/17/2023 | 2404 | 0.939 | 2258 | 123 | 10 | 13 |
| flu B GenBank | before 3/18/2023 | 1000 | 0.996 | 996 | 0 | 4 | 0 |
|  | after 3/17/2023 | 1000 | 1.000 | 1000 | 0 | 0 | 0 |
| flu B EMBL+DDBJ | before 3/18/2023 | 391 | 0.972 | 380 | 1 | 0 | 10 |
|  | after 3/17/2023 | 99 | 0.990 | 98 | 0 | 0 | 1 |
| flu C GenBank | before 3/18/2023 | 16 | 0.875 | 14 | 0 | 0 | 2 |
|  | after 3/17/2023 | 15 | 1.000 | 15 | 0 | 0 | 0 |
| flu C EMBL+DDBJ | before 3/18/2023 | 500 | 0.732 | 366 | 1 | 0 | 133 |
|  | after 3/17/2023 | 130 | 0.400 | 52 | 0 | 0 | 78 |
| total | before 3/18/2023 | 21907 | 0.980 | 21464 | 208 | 81 | 154 |
|  | after 3/17/2023 | 13648 | 0.982 | 13406 | 123 | 27 | 92 |

### Defining pass/fail outcomes

To compare FLAN and VADR validation and annotation using our training and testing sets, we examined each program’s pass or fail outcome for each sequence. For VADR, a sequence fails if one or more fatal alerts are reported for it and passes otherwise. For FLAN, we defined a sequence as failing if one or more errors, denoted with a line starting with “ERROR” in the output feature table and listed in Table 2, are reported, with four exceptions. First, we ignored any errors for PB1-F2 (influenza A segment 2), PA-X (influenza A segment 3), or NB (influenza B segment 6) CDS features because Genbank curators routinely permit problems in these coding regions, annotating them as “misc feature”s instead of CDS. This is because NB is nonessential for virus replication [11], PB1-F2 appears to be nonessential for viral viability [7], and PA-X has variable stop positions which makes validation problematic [13]. Ignoring errors for these CDS is consistent with our VADR tests, for which most fatal alerts in these CDS do not cause a sequence to fail due to a special setting in the VADR model files. Secondly, FLAN frameshift errors for mature peptides were ignored because every instance was accompanied by a frameshift error in the parent CDS (always the HA CDS), and because VADR only reports CDS frameshifts. Thirdly, the error “Input sequence is too short” was reported for 12 sequences in our training and testing sets of 200 nt or less (minimum length 61 nt (FJ222309.1) and maximum length 200 nt (FR687037.1)), however, there were 68 other sequences of 200 nt or less that FLAN did not report this error for (e.g. AM922136.1, 63nt), and VADR does not have an analogous alert, so we decided to ignore this error. Finally, FLAN reported an error with the explanation: “Expected splice site consensus sequence not found for protein 2” for four total sequences (e.g HE584752.1) in our training and testing sets in which only one protein/CDS was annotated. Because it was not clear which protein the error referred to, we decided to ignore these four error instances.

## Results and discussion

The VADR models built from the FLAN reference sequences were used to annotate the training sequence sets using the *v-annotate*.*pl* program, and the VADR results were compared with FLAN annotation results for the same sequences. Differences were manually examined to identify deficiencies in the VADR models, in the form of failing sequences that should pass, which were addressed by adding eight additional influenza A proteins to the VADR *blastx* libraries. For the CY002284 segment 8 model, three additional NS1 (nonstructural protein 1) protein sequences were added (from nucleotide accessions KT370023.1, MT261562.1 and MT169392.1); for the CY002010 and CY002538 segment 6 models, the OP775156.1 and KT181405.1 NA (neuraminidase) proteins were added, respectively; for the CY004642 segment 4 model, the JF715039.1 HA (hemagglutinin) protein was added; for the CY003646 segment 2 model, the AB586849.1 PB1 (polymerase PB1) protein was added; for the CY003645 segment 3 model, the LC625435.1 PA (polymerase PA) protein was added.

Two significant modifications were made to the VADR software to better replicate the FLAN results. The first was the ability to detect and report extra sequence at the 5’ and 3’ ends of sequences with the *extrant5* and *extrant3* VADR alerts, analogous to the FLAN errors with the description “contains extra X nts upstream/downstream the consensus 5’/3’ end sequence.” The new *extrant5* and *extrant3* alerts are similar to the preexisting *lowsim5s* and *lowsim3s* alerts for low similarity at the 5’ and 3’ ends respectively, but *extrant5* and *extrant3* specifically pertain to sequence that extends past the 5’ and 3’ termini of the reference model based on an alignment to the model, whereas *lowsim5s* and *lowsim3s* are reported when the ends of the input sequence do not match well to the model, regardless of whether the sequence alignment extends past the end of the model or ends internally as it would for partial genome sequences.

The second modification is the detection of canonical GT/AG donor/acceptor splice sites at the ends of intron sequences for the PA-X (A segment 3), M2 (A segment 7), M1 (C segment 6), and NEP (A and B segment 8, C and D segment 7) genes, and the reporting of *mutspst5* and *mutspst3* alerts when those expected subsequences are missing in the input sequence. These new alerts are analogous to FLAN’s “Expected splice site consensus sequence not found for protein X” errors. This capability did not yet exist because none of the other viruses or genes that VADR has been used for previously at NCBI include introns.

Table 5 summarizes the results of VADR and FLAN on the training sequence datasets. To demonstrate the impact of different aspects of the models, results for three different VADR model sets are shown: “FLAN-nt” are VADR models built exclusively from the FLAN reference nucleotide genomes and proteins from those genomes only; “FLAN” models additionally include the full FLAN reference protein sets, and “final” are the final set of VADR models. The final models include the eight additional proteins not present in the FLAN reference datasets specifically added to allow additional high quality influenza A training sequences that would otherwise fail to pass VADR. The final set also include the 29 alternative and novel VADR models not present in FLAN, as detailed in Materials and Methods.

Using the final models, VADR and FLAN give the same pass/fail designation for 97.8% or more of the sequences in each of the influenza A and B training datasets, indicating that VADR nearly always reproduces the pass/fail determination of FLAN for A and B sequences. Two hundred and ten type A and four type B sequences fail both programs. For influenza C, nearly all sequences either pass both programs or fail FLAN but pass VADR. In the set that fail FLAN but pass VADR, all but four of the 230 sequences fail FLAN due to extra sequence at the 5’ or 3’ end which are not reported by VADR due to the alternative, longer influenza C models it uses. We did not test influenza D because FLAN does not include influenza D reference sequences.

The addition of the FLAN protein sets to the nucleotide only models (“FLAN” vs “FLAN-ntonly”) results in a large increase in the number of influenza A sequences that pass VADR, demonstrating the importance of multiple reference proteins for at least some of the influenza proteins. In the influenza A GenBank and non-GenBank sets (10,000 sequences each) 908 and 496 sequences, respectively, that failed VADR using the “FLAN-ntonly” models pass with the “FLAN” models. The corresponding increase in the influenza B datasets (1,000 sequences each) is 55 in the GenBank set and 7 in the non-GenBank set. For influenza C these is no change.

Supplementing the “FLAN” influenza A models with additional proteins to the VADR protein libraries which are not present in the FLAN sets (“FLAN” vs “final”) results in a smaller increase in the number of influenza A sequences that pass VADR. Specifically, 32 and 58 sequences that previously failed VADR but passed FLAN now pass both programs in the GenBank and non-GenBank sets respectively.

We also ran VADR and FLAN on our testing sets, which are disjoint from the training sets, to check how general the improvements made based on our analysis of the training set results would be on other sequences. The “final” model sets were already defined prior to our examination of the test set results, and no further changes were made based on their analysis. The test set results largely mirror the training set results, as shown in Table 6, which only shows results for the final model set, demonstrating that the models are likely not overtrained on the training data and should yield comparable results on new sequences. The high consistency between FLAN and VADR results on the new sequences collected since March 17, 2023 also suggests VADR performance on sequences outside of the training set should largely be consistent with FLAN (Table 6).

### Sequences with different VADR and FLAN pass/fail outcomes

We manually examined each sequence in the training and testing sets for which VADR and FLAN give different pass/fail outcomes. Below, we summarize our findings separately for each influenza type (A, B or C).

### Differences for type A sequences

There are 204 influenza A sequences in the training and testing sets that pass FLAN but fail VADR. Eighty five of these fail due to a possible frameshift (*fsthicft* and *fsthicfi* VADR alert codes), with nearly half (40) in the hemagglutinin CDS. A typical example is shown in Figure 1. Of these 85, 77 are terminal frameshifts near the beginning or end of the sequence, and many are short frameshifts: 70 are 15 nt or less. Of the remaining 119 sequences, 44 fail VADR protein validation because no *blastx* alignment extends sufficiently close to the 5’ and 3’ ends of the predicted CDS (*indf5pst* and *indf3pst*, e.g. CY008965); 40 fail due to extra sequence detected at the 5’ or 3’ ends relative to the reference model sequence (*extrant5, extrant3* alerts, e.g. OR675339.1); 11 fail due to large deletions (*deletinp* alerts, e.g. OY283585.1); eight fail due to possible mutations at splice donor or acceptor sites (*mutspst5, mutspst3*, e.g. AB513916.1); seven fail with low sequence similarity to the reference at the 5’ and/or 3’ ends of the sequence (*lowsim5s, lowsim3s*, e.g. OQ683476.1); five fail due to possible mutations in the stop codon (*mutendex, mutendcd, mutendns*, e.g. LC070034.1); one fails due to a mutation in a start codon (*mutstart*, e.g. MH283613.1), one fails due to low coverage resulting from a large stretch of N ambiguity characters of unexpected length (*lowcovrg*, CY095556.1), one fails due to a large insertion (*insertnp*, OQ722117.1), and one fails due to a large deletion (*deletinp*, OY282827.1).

**Fig. 1:**
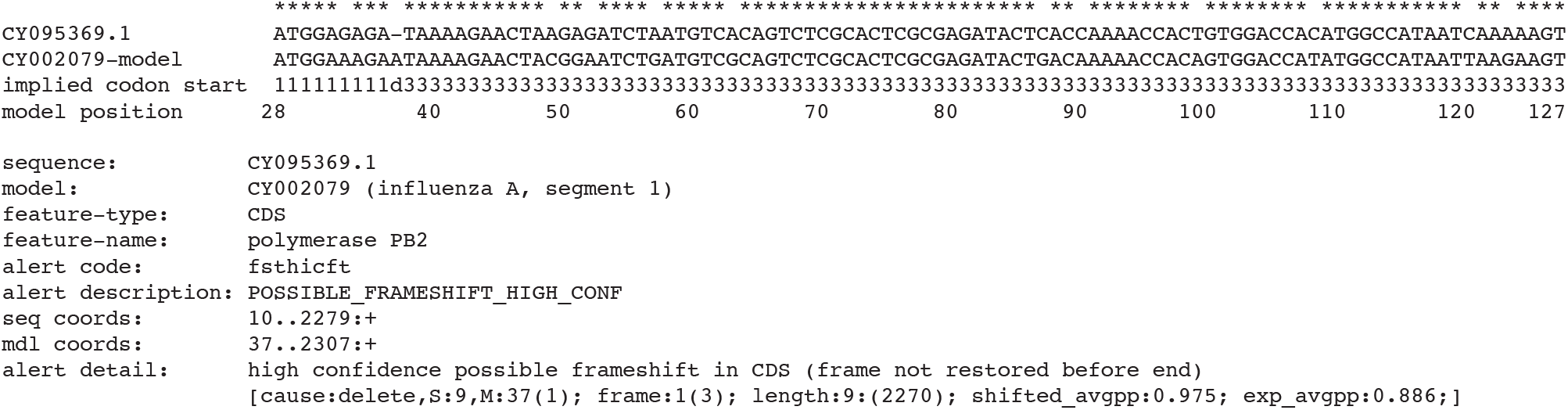
Example potential frameshift detected by VADR but not by FLAN. VADR alignment of the first 99 nucleotides of CY009539.1 to the CY002079 influenza A segment 1 model reference sequence is shown with a single deletion with respect to the model sequence at model position 37. The polymerase PB2 CDS is encoded by positions 28 to 2307 of CY002079 so the first three nucleotides of the alignment correspond to the start codon. VADR reports a potential frameshift (*fsthicft* alert) of all nucleotides (positions 10 to 2279) after the deletion. Identical aligned nucleotides between the sequence and the model are indicated by * at the top of the alignment. Some of the information reported in the VADR output file with suffix “.alt” is included below the alignment. FLAN passes CY002079 without a frameshift error or any other errors, possibly because the 9 nucleotide length prior to the frameshift is so short.

For some of the sequences that fail due to extra sequence alerts (*extrant5* or *extrant3*), the additional sequence is actually a duplication of a sometimes reverse complemented region of the genome (e.g. LC168638.1), possibly due to an assembly error, which VADR reports as a separate *dupregin* and/or *indfstrn* alert. For some of the eight failures related to mutations at splice sites (*mutspst5, mutspst3*) the sequence fails because the intron for the M2 gene has been removed (e.g. EU384412.1, which matches best to the CY002009 model for segment 7).

We have manually inspected all of these VADR failures and believe that they are warranted in nearly all cases because they are situations that GenBank curators should manually investigate prior to the deposition of these sequences in GenBank. In a few cases VADR simply gets it wrong and the sequences could reasonably pass without manual examination. An example is LC120391.1 which involves the deletion of a single G relative to the reference sequence CY000449, the first G in the subsequence TGAG within which TGA is the stop codon for the hemagglutinin CDS, but which leaves a valid TAG stop codon. VADR’s alignment-based stop-codon annotation balks at this deletion and reports a MUTATION_AT_END error (*mutendex, mutendcd* and *unexleng* alerts). This situation, which is fortunately rare in influenza sequences, highlights a specific limitation of VADR.

There are 37 influenza A sequences that pass VADR but fail FLAN in our training and testing sets. Twenty of these fail due to ambiguous nucleotides in the start and/or stop codons of CDS, which VADR is tolerant of (e.g. OX422492.1). For 11 sequences FLAN reports a mutation in the start or stop codon (e.g. EU146784.1), but in all cases there is a valid start/stop codon that corresponds to at least one protein in the FLAN reference set (which is why these sequences do not fail VADR). A “wrong exon number” error is reported by FLAN for three sequences with more than 500 Ns (e.g OY284219.1). VADR’s method of replacing large stretches of Ns with reference sequence for validation and annotation purposes, and which does not have an analog in FLAN, seems to work well for these N-rich sequences. FLAN reports that two of the remaining three sequences (FJ222261.1 and FJ222309.1), both 61nt, do not have coding capacity, even though they encode 20 amino acid long partial NEP CDS. The final sequence (GU083607.1) fails due to extra nuleotides upstream of the consensus 5’ end sequence.

We have manually inspected these 37 sequences and have determined that in our opinion they should all pass. This does not mean that it is a significant problem that FLAN fails these sequences, as it is reasonable for some borderline sequences to fail so that expert curators can evalutate them before they are deposited into GenBank. However, it is reassuring to find that VADR is not allowing sequences to pass that are clearly problematic.

### Differences for type B sequences

In the combined influenza B training and testing sets, there are 26 sequences that pass VADR but fail FLAN. For 17 of these FLAN reports a probable mutation at start of the NEP gene (nonstructural protein 2), but in all cases the sequences begin towards the end of the first exon of NEP (e.g. AJ781277.1) or within its intron (e.g. AJ781285.1). Of the remaining nine sequences, eight are partial segment 7 sequences that end before the start of the BM2 CDS (e.g. HE803092.1) but FLAN reports a probable mutation at the start of the BM2 CDS. The final sequence (OY759012.1) fails due to probable mutation at start and end due to ambiguities in the start and stop codons which VADR is tolerant of. We manually examined the VADR annotation for these 26 sequences and found it to be correct and confirmed that in our opinion none of the errors were significant enough to prevent deposition in GenBank.

Of the five influenza B training and testing sequences that fail VADR but pass FLAN, four are due to possible frameshifts in segment 4 (e.g. KP461008.1) and one is due to 11 extra nucleotides at the 3’ end (MN086295.1). Three of the four frameshifts are due to a single nucleotide deletion at reference position 85 of the model sequence AF387493.

### Differences for type C sequences

For type C sequences, there are 443 sequences in the training and testing sets that pass VADR and fail FLAN. The vast majority of these (429) fail only due to FLAN errors about extra nucleotides upstream or downstream of the consensus 5’ or 3’ end sequence owing to extra sequence on the 5’ and/or 3’ ends (e.g. AF170575.2) beyond the FLAN influenza C reference sequences. VADR uses longer influenza C RefSeq sequences of which the FLAN references are subsequences (see Materials and Methods) which result in zero VADR *extrant5* or *extrant3* errors for these 429 sequences. The other 14 sequences fail FLAN due to probable mutation at end errors which upon manual inspection appear to be invalid errors as the sequences all clearly end prior to the end of the CDS the error is reported for (e.g. the CM2 CDS for D78423.1).

### Error comparison

Table 2 lists all FLAN errors, corresponding VADR alerts and error messages and counts in the combined training and testing sets. FLAN reports 1297 of these errors and VADR reports 1008 of its analogs of those errors. Of these, 642 are consistent in that the same error is reported for the same feature (CDS or mature peptide), but 655 FLAN errors and 366 VADR errors are unique (not consistent). The majority of these unique error instances were examined and listed in our analysis of the sequences with different pass/fail outcomes above. For example, at least 429 of the 655 unique FLAN errors were influenza C sequences that failed due to extra nucleotides upstream or downstream of the 5’ and/or 3’ ends (“at least” because one sequence could have both errors).

### Classification and annotation comparison

Of the 58,555 total sequences in our training and testing sets, FLAN and VADR classify all but three to the same type, segment and subtype (for type A segments 4 and 6). FN395357.1, AM922160.1, and ON527769.1 are classified by FLAN as H2, H4, and H16 subtypes, respectively, but as H1, H10, and H1 by VADR. In all three cases, the top *blastn* hits against “nr” support the VADR classification.

There are 196,683 total features annotated in our training and testing sets by one or both programs. Of these, both programs give identical start and stop coordinates for 194,119 (98.7%). There are 1331 (0.7%) features annotated by both programs but with different coordinates, 827 (0.4%) features that VADR annotates that FLAN does not, and 406 (0.2%) features that FLAN annotates that VADR does not.

Taken together, these data demonstrate that VADR, when employing models derived mainly from FLAN’s reference nucleotide and protein sequences, largely reproduces FLAN’s pass/fail, classification and annotation results. The majority of problems detected by either program are detected by both, and analogous alerts or errors are reported. When the two programs differ, VADR nearly always provides the more appropriate pass/fail decision.

### Efficacy of the novel VADR models

As noted above, the VADR influenza model set (v1.6.3-2) contains 17 additional influenza A, seven additional influenza D and five alternative influenza C models that are not derived from FLAN reference sequences. While there are very few INSDC sequences that match best to the 17 additional influenza A models based on novel H and N subtypes at the time of writing, using these additional models does improve VADR performance on those few sequences.

We used VADR to annotate all 977,136 influenza A sequences listed in NCBI virus on December 13, 2023 and found that only 78 match best to one of the 17 added models for H17/N10, H18/N11 or H19 subtypes. Of these, 57 were classified to models that matched their annotation in GenBank (e.g. KR077935.1 matched best to the CY125948 model and is annotated in GenBank as subtype H18/N11). All 57 of these passed, and only 13 would have passed if the 17 new models were not added. For the other 21 sequences, VADR almost certainly misclassified them to the new subtypes, in many cases probably due to their short length which can make classification more challenging (19 of the 21 are 90 nt or shorter). However, only four of these 21 failed VADR, and this is the identical set of four that would fail if run against the set of models without the new models. The new influenza C models also improve performance. As mentioned above, using the five new longer influenza C RefSeq models instead of models based on the non-RefSeq FLAN reference sequences allows 429 sequences in our training and testing sets to pass VADR that fail FLAN due to extra nucleotides upstream or downstream of the consensus end sequence.

### Limitations and future directions

VADR is designed to be a general tool capable of validating and annotating most viruses with genomes less than about 30Kb. While building models using the *v-build*.*pl* module is straightforward, we have found that significant testing and manual modification of viral models is often necessary to achieve the level of accuracy and reliability necessary to use them in an automated fashion, such as in the context of automatically screening incoming GenBank sequence submissions. Our experience with influenza reinforces this as we were able to significantly improve performance of the initial models built from only the FLAN nucleotide reference sequences by adding additional proteins to our *blastx* libraries (mostly from FLAN’s set, see Table 5, “FLAN-nt” vs “final” rows). The preexistence of FLAN’s reference protein sets made this task much easier than it would have been otherwise. This required testing and model improvement is currently a major bottleneck in creating sufficiently accurate VADR models.

The relatively slow speed of VADR in annotating sequences can be limiting in some contexts. VADR is especially slow for longer sequences due to the high complexity of its search and alignment algorithms, although this can be alleviated for viruses with relatively low sequence variability (e.g. SARS-CoV-2 [18]). For influenza sequences, which are nearly always less than 2500 nt, speed is less of an issue, especially for typical input datasets of less than 1000 sequences. When parallelized across 16 processors (using the `split --cpu 16` options), *v-annotate*.*pl* processed all 977,136 influenza A sequences in INSDC (listed in the NCBI virus resource on December 13, 2023) in just under 24 hours on 2.2 GHz Intel Xeon processors. This averages to about 2500 sequences per processor per hour, or about 1.5 seconds per sequence per processor. This speed is in the same ballpark as FLAN, which took about 14 minutes to annotate 1000 influenza A sequences from our training set in our hands, or about 0.85 seconds per sequence, making it about twice as fast as single-processor VADR. Our recommended usage of *v-annotate*.*pl*, detailed in the documentation included with the influenza models, employs four processors which ends up making VADR about twice as fast as FLAN in practice.

An important feature of FLAN is its ability to detect frameshifts using the ProSplign DNA to protein alignment software. ProSplign computes alignments at the protein level, by translating the input sequence and aligning it to reference proteins. VADR also attempts to detects frameshifts but does so at the nucleotide level, aligning the input nucleotide sequence to a reference genome sequence. In principle, when comparing sequences working in protein space is more powerful due to the larger alphabet and higher information content of protein sequences [20]. However, our testing of influenza annotation suggests that this distinction does not confer a large advantage to FLAN. We examined all of the sequences that fail due to frameshifts in FLAN but not VADR, and in each case the frameshift call looked incorrect. Working in nucleotide space actually seems to allow VADR to detect potential short frameshifts that FLAN does not report. An example is shown in Figure 1.

Using VADR should enable simple model changes in the future. As novel influenza diversity is observed in INSDC sequence data, the VADR development team will work together with the NCBI Virus team to identify reference sequences for novel proteins or subtypes and use them to update the VADR models. While in principle FLAN models are also expandable in this way, in practice new reference sequences and proteins were rarely added.

## Conclusions

The amount of viral sequence data generated by independent research and public health labs and submitted to public sequence databases will likely continue to grow. This underscores the need for publicly available tools that can be easily maintained and updated as new sequence diversity is encountered. The VADR package is a general tool that allows the validation and annotation of different types of viruses including several flaviviruses, caliciviruses, coronaviruses and now influenza (*Orthomyxoviridae*) using a single codebase. Since 2007, GenBank has used the FLAN software to validate and annotate incoming influenza sequences. Using models derived from the reference sequences used by FLAN, VADR gives very comparable annotation to FLAN on influenza A, B, and C sequences, and additionally allows annotation of influenza D sequences. When there are discrepancies, VADR usually gives more accurate annotation. Based on our findings, we intend to start using VADR instead of FLAN for GenBank influenza sequence processing.

## Funding

This research was supported by the Intramural Research Program of the National Institutes of Health, National Library of Medicine (NLM), and by the National Center for Biotechnology Information of the NLM. As a funding body, the NLM had no role in the design of the study, the collection, analysis, and interpretation of the data, and no role in writing the manuscript.

## Conflict of interest

None declared.

## Data availability

VADR and the influenza models are in the public domain and are freely available at https://github.com/ncbi/vadr and https://bitbucket.org/nawrockie/vadr-models-flu. VADR depends on the following software, which is downloaded and installed as part of VADR installation: Bio-Easel v0.16, BLAST+ v2.15.0, Infernal v1.1.5, FASTA v36.3.8h, minimap v2.26 and Sequip v0.10. All data generated or analyzed during this study are included in this published article, its supplementary material, or NCBI’s GenBank database. The VADR influenza models dataset (vadr-models-flu-1.6.3-2.tar.gz) includes information on mapping the FLAN reference sequences to INSDC sequences. The supplementary material includes information on the the collection and analysis of the training and testing datasets as well as instructions for reproducing the tables in the paper (https://github.com/nawrockie/vadr-flu-paper-supplementary-material).

## Acknowledgements

We thank the NCBI systems team, especially Ron Patterson, for management of the compute farm.

## Notes

https://github.com/nawrockie/vadr-flu-paper-supplementary-material

### Competing Interest Statement

The authors have declared no competing interest.

